# Future climate change promotes novel gene-climate associations in balsam poplar (*Populus balsamifera* L.), a forest tree species

**DOI:** 10.1101/2020.02.28.961060

**Authors:** Andrew V. Gougherty, Stephen R. Keller, Vikram E. Chhatre, Matthew C. Fitzpatrick

## Abstract

A central challenge to predicting climate change effects on biodiversity is integrating information on intraspecific variation, specifically population-level local adaptation to climate. Assessing how climate change could disrupt local adaptation to climate can provide a new way of understanding population risk and vulnerability to climate change. For the wide-ranging boreal tree species, balsam poplar (*Populus balsamifera* L.), we used models of existing population-level genetic differentiation to estimate three key components of population’s vulnerability to climate change: (1) predicted shifts in genetic composition with and without migration, (2) the potential for future novel gene-climate associations, and (3) the distance populations would need to migrate to minimize future maladaptation. When assessed across the range of balsam poplar, these three metrics suggest that vulnerability to climate change is greatest in the eastern portion of balsam poplar’s range, where future maladaptation peaked, migration distances to sites that minimized maladaptation were greatest, and the emergence of novel gene-climate associations were highest. Our results further suggest greater maladaptation to climate when migration distances were limited – consistent with the possibility of migration to lessen maladaptation to future climate. Our work provides a comprehensive evaluation of population’s vulnerability to climate change by simultaneously assessing population maladaptation to future climate and the distances populations would need to migrate to minimize maladaptation, in a way that goes beyond species-level bioclimatic modelling. In doing so, our work helps advance towards the long-held goal of incorporating genomic information in models of species responses to climate change.

## MAIN TEXT

To understand impacts of climate change on climatically-adapted populations there is a need to simultaneously assess the potential for populations to become maladapted to future climates, and the distance populations may need to migrate to minimize future maladaptation^1,2^. Most efforts to quantify species’ responses to climate change, however, take a species-level approach that assumes all populations within the range respond similarly to environmental change, and ignore the potential role of local adaptation to climate. While techniques have been developed to integrate intraspecific variation with predictions of species range shifts^3–5^, they rarely capture the continuous nature of adaptive variation present across species’ ranges. Moving beyond species-level approaches, such as species distribution models, to directly predict climate change effects on populations has the potential to provide new, nuanced insights into population vulnerability to climate change that may help inform where conservation efforts may be most effective at preventing extirpation of vulnerable populations.

Climatically-adapted populations are likely to face numerous challenges from shifting climates. Disappearing and novel climates^6^, for instance, could be particularly troublesome for climatically-adapted populations if they result in populations becoming maladapted to climate. Populations adapted to disappearing climates (i.e., contemporary combinations of temperature and precipitation with no future analogs) may be extirpated from the landscape in the absence of genetic rescue by geneflow or allele frequency change in situ. In turn, population extirpation will likely result in the loss of rare alleles within the range^7^ and could reduce the adaptive potential of populations to future stresses. Alternatively, novel climates, which existing genotypes have no prior selection history, could favor the evolution of novel gene-climate associations through the process of recombination of existing climate-adaptive alleles. High climate novelty could cause range retractions if future climates in the range are far outside the current climatic tolerances of existing populations. While both disappearing and novel climates are likely to be detrimental to populations *in situ*, migration has the potential to dampen the effects. If populations can disperse quickly (or be moved) to areas with climates they are preadapted to, the risk of extirpation and range retraction could be lessened. Absent natural migration, moving individuals long distances has been proposed as a way to conserve vulnerable populations by minimizing future maladaptation. Quantifying the relative need for populations to adapt and migrate, hence, can provide a direct assessment of population’s risk to future climate change.

Here, we integrate concepts of novel and disappearing climates with adaptive genetic variation to map where within species’ ranges populations may be most preadapted or maladapted to future climates, and the distances populations would have to migrate to minimize future maladaptation (Fig. 1). As a case study, we use a subset of locally adapted genes associated with phenology in balsam poplar (*Populus balsamifera* L.), a wide-ranging northern North American tree species. Using generalized dissimilarity models (GDM), we modelled differentiation of adaptive genetic variation (measured as *F*_*ST*_) as a function of climatic differences between 81 populations, and calculated three metrics of genetic offset: local, forward, and reverse offset. Genetic offset, following Fitzpatrick & Keller^8^, represents the disruption of the current gene-climate relationship due to shifts in climate. Local offset assumes populations do not migrate in response to climate change and, hence, reflects only the effects of local shifts in climate (Fig 1.2a). Local offset was calculated by predicting *F*_*ST*_ due to local changes in climate in balsam poplar’s range over the next fifty years (2070). Forward offset, on the other hand, assumes populations have unlimited migration ability and was calculated by identifying the minimum predicted *F*_*ST*_ if populations could migrate/be moved anywhere in North America (Fig. 1.2b). Forward offset can be interpreted as the relative possibility of contemporary gene-climate associations disappearing from the landscape if populations were unconstrained by migration. Reverse offset is similar to the concepts of novel climate or novel species assemblages^9,10^, but applied to the adaptive genetic composition of populations - representing the adaptive novelty required of current populations to be preadapted to future climates. Reverse offset was calculated by identifying the minimum *F*_*ST*_ between hypothetical populations in future climate and populations in current climate (Fig. 1.2c). We also mapped the distance and direction populations would migrate to minimize forward offset, and tested the effect of limiting migration to five distance bins (50, 100, 250, 500, 1000 km) on forward offset.

**Fig. 1.**
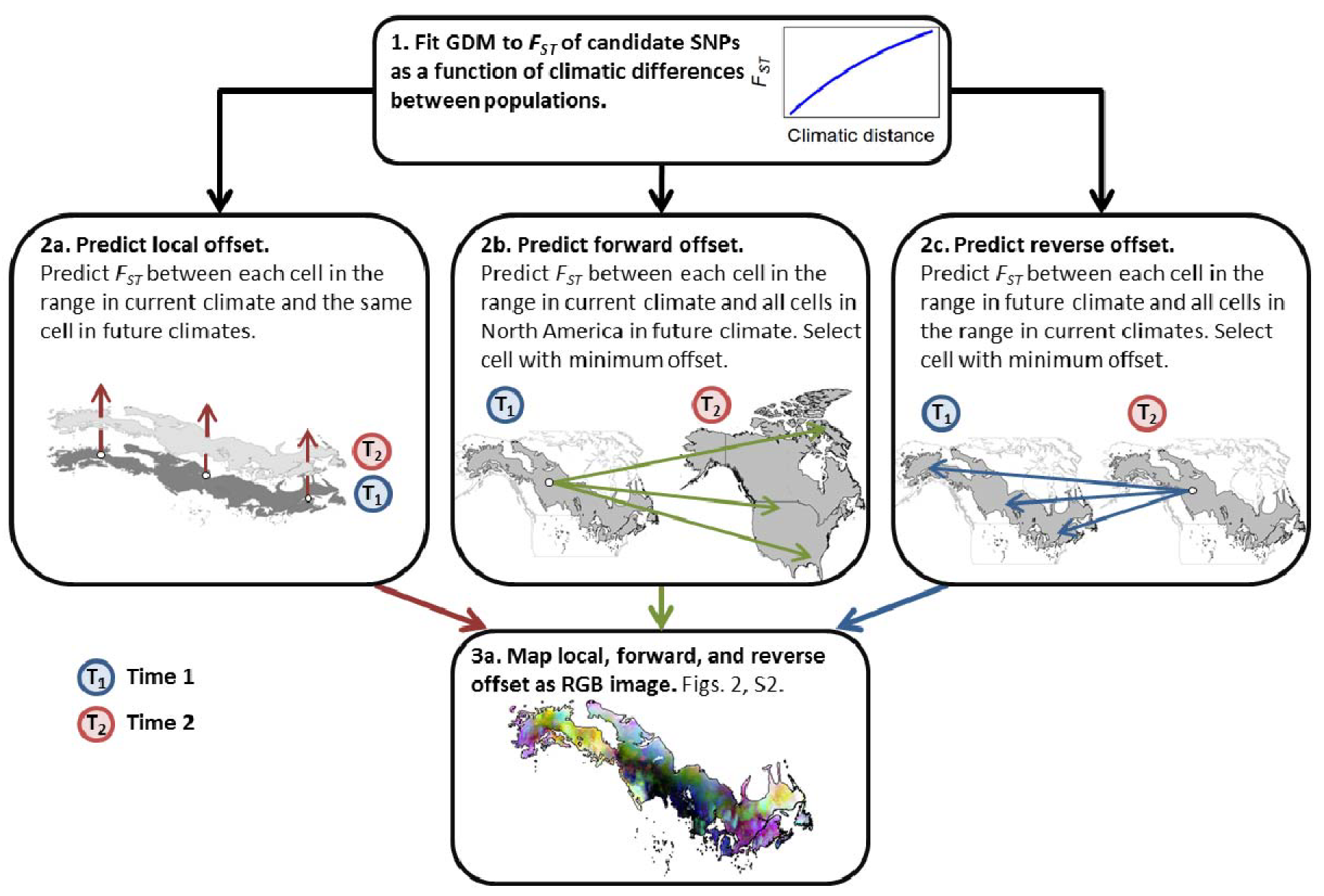
Schematic of how local, forward and reverse offset were calculated and mapped. (1) After fitting a generalized dissimilarity model to *F*_*ST*_ of climatically-adaptive SNPs, the model is used to predict (2a) local, (2b) forward, and (2c) reverse offset. Local offset is calculated following Fitzpatrick & Keller ^8^. Forward offset is calculated by predicting *F*_*ST*_ between each cell in the range in current climate and all cells in North America in future climate and selecting the minimum value. Reverse offset is calculated by predicting *F*_*ST*_ between each cell in the range in future climate and all cells in the range in current climate and selecting the minimum value. Gray polygons are balsam poplar’s range ^22^.

**Fig. 2.**
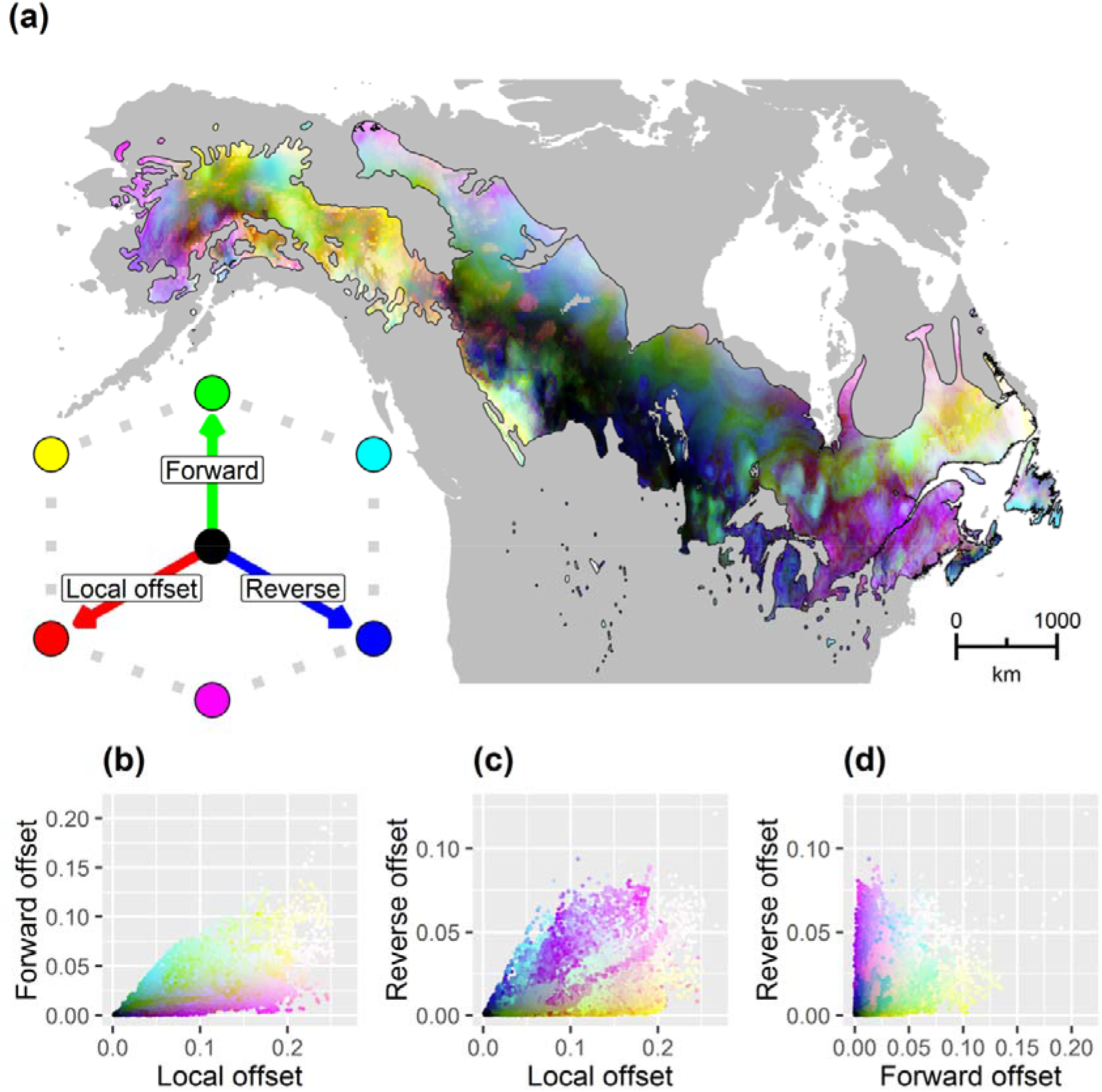
Red-green-blue map of local (red), forward (green), and reverse (blue) offset throughout the range of balsam poplar for 2070 and RCP 8.5. Brighter cells (closer to white) have relatively high values along each of the three axes indicating greater predicted exposure to climate change, while darker cells (closer to black) have relatively lower values, indicating lower exposure to climate change. (b-d) Bivariate scattergrams of (a). All units are *F*_*ST*_, but forward and reverse offsets are minimized *F*_*ST*_.

While the predicted patterns of local, forward, and reverse offset were highly variable within balsam poplar’s range, some generalities did emerge (Fig. 2, S2, S4). Local, forward, and reverse offsets were each lowest in the center of the range. Populations in the center of the range were predicted to (i) have low offset in their current location (i.e., with no migration), (ii) have low offset in other regions of North America (i.e., could migrate somewhere in North America with low adaptive offset; low forward offset), and (iii) existing populations elsewhere in the range have low predicted offset for future climate in the center of the range (i.e., low reverse offset). Together, low local, forward, and reverse offsets suggest populations in the center of the range may be the most preadapted to future climate for the loci we assessed. In contrast, the easternmost and northernmost parts of the range were predicted to have high local, forward, and reverse adaptive offsets (Fig. 2). This indicates there are no existing balsam poplar populations, either locally or elsewhere in the range, that are predicted to be preadapted to future climates in the eastern and northern parts of the range. High local and forward offsets suggest eastern and northern populations are likely to be particularly vulnerable to climate change as the impacts of local climate shifts cannot be mitigated by migration or movement of populations to regions with more suitable climate. As such, populations in the eastern and northern parts of the range are likely to be most at risk of extirpation due to climate change, potentially resulting in range contraction near the contemporary longitudinal range edges.

### Migration distances and direction

Interestingly, migration distances (i.e., the distance to locations that minimize forward offset) were only weakly correlated with forward offset (*r* = 0.14, *p* = 0.25) – suggesting that, across populations, the need to migrate longer distances was not necessarily associated with a higher adaptive offset. Distances to locations that minimized future maladaptation (*D*_*min*_) peaked in the eastern portion of the range, where they exceeded 5000 km (Fig. 3a). *D*_*min*_ for many of the cells in the northeastern portion of the range were in mountainous regions in the western half of North America – indicating populations in the easternmost portion of the range would need to migrate (or be moved) across nearly the entire North American continent to minimize future maladaptation. The shortest *D*_*min*_ values, in contrast, occurred along the southern range edge, and sporadically in the northern portion of the range, often near mountainous areas. Very few locations (<1%) had *D*_*min*_ of zero, suggesting that populations in nearly all parts of the range would need to migrate some distance to reach the location they are most preadapted to in the future, barring allele frequency change in situ. Limiting the maximum allowable migration distance (i.e., search distance) for forward offset revealed a negative relationship between forward offset and distance (Figs. 4, S5, S6), suggestive of a tradeoff between migration distances and forward offset. Forward offset decreased considerably when the search distance was expanded from 0 to 500 km (mean decrease in forward offset across all cells: 53.9%), while the decline in forward offset between distances of 500 and 1000 km was considerably lower (mean decrease: 14.9%), indicating a declining benefit of lowering forward offset as search distances increased. The relative range-wide pattern of forward offset was similar across all search distances but the magnitude differed (Figs. S5, S6).

**Fig. 3.**
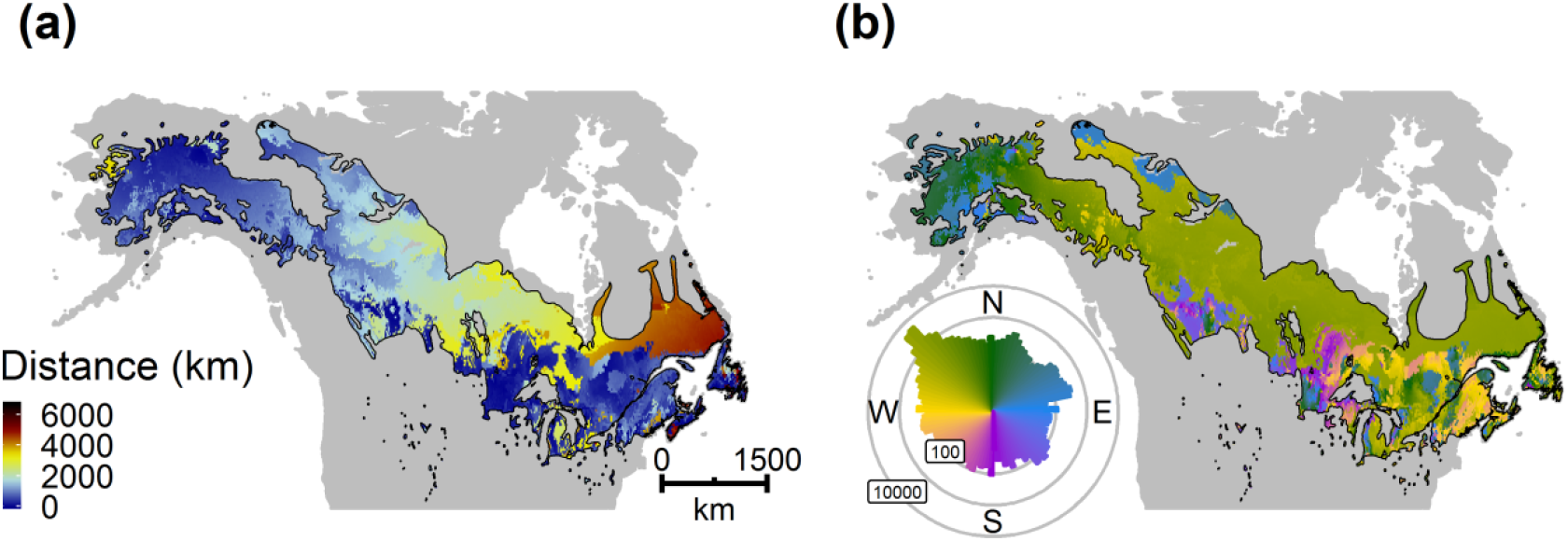
(a) Distance and (b) initial bearing to the location that minimizes future offset for balsam poplar in 2070 and RCP 8.5. Polar histogram in (b) shows the log10 number of cells in each bearing bin.

**Fig. 4.**
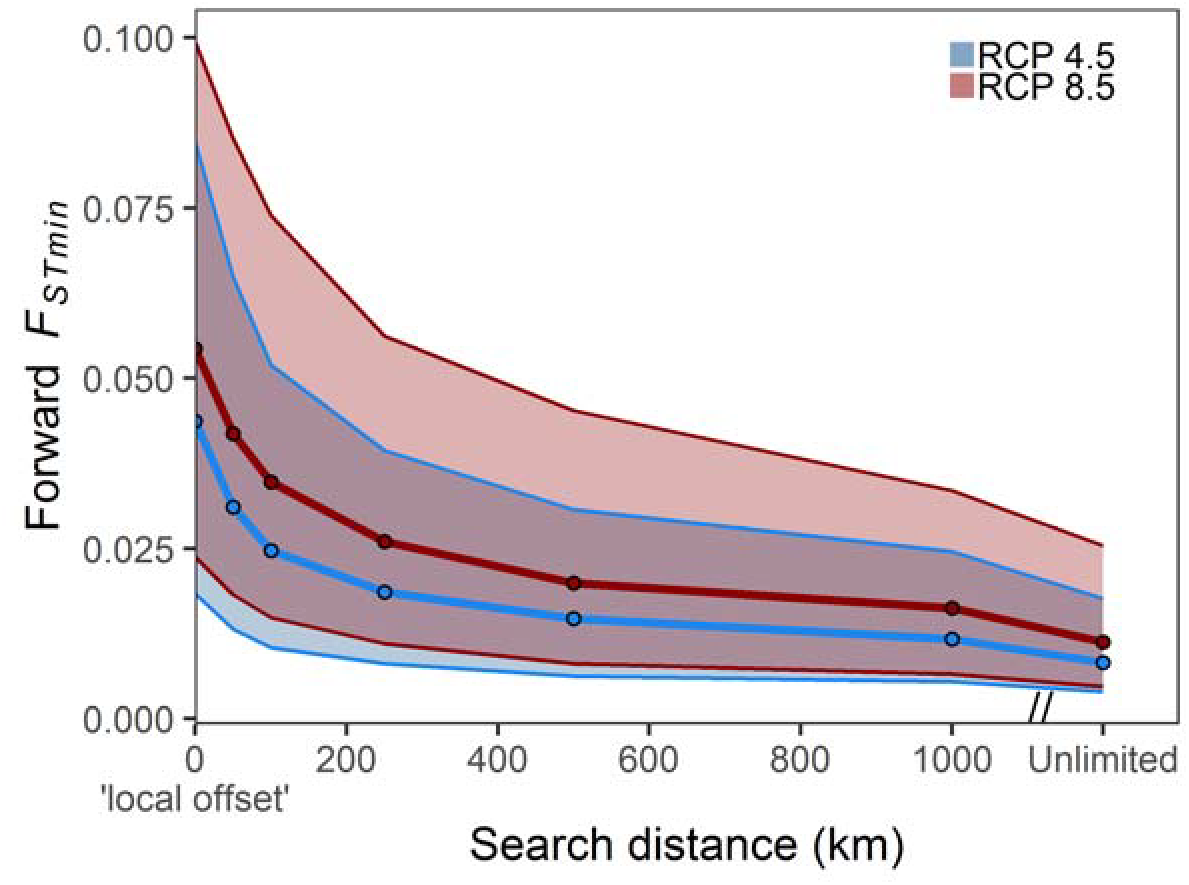
Relationship between maximum allowable migration distance and minimized forward offset (*F*_*STmin*_) for 2070. Bands extend between the 25th and 75th percentiles, and points are median values. See also Fig. S5 & S6.

Our analyses further indicate populations may not migrate strictly northward to minimize future maladaptation. While species-level bioclimatic models typically predict that species will shift their ranges poleward in response to warming temperatures, our population-level perspective reveals that if populations migrate towards the location that minimizes forward offset, range shifts may be considerably more complex. For example, while most (82.8%) locations in balsam poplar’s range had an overall northward trajectory (i.e., the cell that minimized forward offset was at a higher latitude than the source cell), we found considerable variability along the southernmost range edge (Fig. 3b). In these southern edge populations, the migration trajectory was sporadically westward, eastward, and even southward. The variability in population-level trajectories was especially apparent in the upper Great Lakes region where directionality varied over short distances. Recent observational studies have shown similar variability when populations are aggregated over entire species^11,12^. Fei et al.^11^, for instance, showed that over the past 30 years, eastern North American tree species have more often shown a westward shift in abundance than a poleward shift. The authors propose this is due to shifting precipitation regimes and moisture availability increasing suitability of eastern tree species in the center of North America. Precipitation variables were similarly among the most important in our GDM (Fig. S1) and likely explain some of the non-northward trajectories in our predictions.

### Advances and applications

Impacts of climate change on populations will be mediated by the collective ability of locally adapted populations to adapt and/or migrate in response. Our study quantifies these varied responses by developing a spatially-explicit understanding of the relative roles of local maladaptation, minimum migration distances, and gene-climate novelty on climatically-adapted populations. In doing so, we attempt to shed new light on several major unresolved questions concerning how climate change will affect climatically adapted populations, specifically: (i) where will climate change cause the greatest mismatch between locally adapted populations and climate?, and (ii) which existing populations may be most preadapted or maladapted to future climates? By simultaneously calculating multiple metrics of maladaptation and incorporating migration distances, our approach provides insight into the tradeoff between *in situ* selection versus migration, and helps elucidate where within the range migration and adaptation may be most effective at reducing the genetic offset under future climates. It is important to emphasize that our approach makes no attempt to assess the actual ability of populations to adapt or migrate, or predict mean fitness and evolutionary response over multiple generations^13^. Nevertheless, our approach provides intuitive metrics of population-level vulnerability to climate change that may serve as a useful baseline for understanding where populations may be most at risk from climate change.

Contrary to some theoretical work (e.g.^14^), we found balsam poplar’s vulnerability to climate change was not greatest along the trailing (southern) edge of the range, but rather at the longitudinal extremes of the range. Part of the reason for this was that the effects of temperature were secondary to the effects of precipitation in driving differentiation of SNPs we investigated. Change in winter precipitation, the most important variable in our model, is predicted to be greatest in the eastern and northernmost parts of the balsam poplar’s range. Greater winter precipitation, combined with warmer winter temperatures, likely result in warmer/wetter winter conditions that populations have not experienced in the recent past, resulting in high adaptive offset in the eastern and northern parts of the range. In the center of the range, and along much of the southern edge, climate shifts are predicted to be relatively modest compared to the eastern portion of the range, leading to relatively lower offset in these areas. These findings are consistent with recent work that has suggested that accounting for local adaptation when predicting range shifts could yield results contrary to the leading-/trailing-edge paradigm of range shifts (i.e., as ranges shift poleward, trailing edge populations are most vulnerable to climate change as they will be the first to experience temperatures outside the species’ climatic niche^15^). Indeed, other modelling studies have shown accounting for local adaptation in climate change predictions can yield unexpected range shifts that are not strictly poleward^16^. Similarly, numerous empirical studies have reported that range shifts in response to recent climate change are rarely uniformly poleward^11,12,17,18^, and may be in multiple directions, including southward. Together, our results point to the possibility that explicitly considering the gene-climate association of multiple loci across multiple climatic gradients could yield a considerably more complex view of population’s responses to climate change than is often implied by a shift poleward in response to increased temperatures.

In addition to providing multiple metrics of population vulnerability to climate change, our approach can potentially be used to inform conservation and restoration efforts. Assisted migration efforts, for instance, which rely on moving individuals to areas with suitable future climates, could benefit from being able to identify which populations are most preadapted to future climate but may not be able to reach those locations via natural dispersal. Dispersal-limited populations are likely to benefit most from assisted migration efforts, and may be the only option for some populations to avoid extirpation. Conversely, populations that are predicted to be maladapted to all future climates could be candidates for targeted germplasm sampling to ensure rare adaptive alleles remain available for future breeding or restoration programs. Genomic offset metrics hence have the possibility of providing a biologically-meaningful metric to guide conservation of climatically-adapted populations, beyond those offered by climate matching approaches or species distribution models.

### Limitations and future work

While our approach offers unique insights into the magnitude of balsam poplar’s responses to climate change, our approach makes numerous simplifying assumptions that may warrant further investigation. First, the offset metrics implicitly assume that the current pattern of genetic differentiation across space is representative of genetic change through time. While any space-for-time substitution may introduce uncertainty in climate change predictions, using SNPs with an *a priori* relationship with climate^19^, related to a temporally/spatially varying trait (i.e., phenology), helps ensure we are modelling a robust, reliable gene-climate signal. Second, our use of a correlative model (GDM) does not account for the genetic forces (i.e., selection, gene flow, etc.) that will shape the future pattern of genetic variation, nor do the models account for potential plastic responses to climate change, or population’s ability to persist in variable climates (i.e., interannual climatic variability) – both of which may contribute to an overestimation of offset metrics. Finally, our models emphasize only adaptation to climate.

While focusing on climate may be suitable for SNPs related to phenology, populations are also likely to experience unique biotic interactions in future climates (e.g., encountering new competitors or new pest regimes). Hence, it is important to emphasize that the offset metrics calculated here are only relevant to the specific SNPs used in our study, and may not be generalizable to other portions of the genome responsive to climate.

While our analyses were conducted on a small subset of loci in balsam poplar associated with phenology, which have a well-studied physiological and phenotypic relationship with climate and is locally adaptive in balsam poplar^20^, our approach is generalizable to any number of loci having a robust adaptive association with climate. Further insights could be gained by assessing the pattern of forward and reverse offset of loci associated with other climatically-adaptive traits (e.g. heat and drought tolerance, growth rates, etc.). Assessing loci associated with other traits could help elucidate the variable impacts climate change may have on different parts of the genome, and could inform whether populations are preadapted to a single location on the landscape, or more likely, whether genomic regions underlying different functional traits will be preadapted to different locations on the future landscape. Such information will be crucial to understanding and mitigating the effects of climate change on local adaptation in forest trees.

## CONCLUSION

Populations, not species, respond to climate change, and these responses are likely to include a complex combination of migration and adaptation to avoid extirpation^1,2,21^. Here, we attempt a first step towards accommodating this complexity by estimating multiple metrics of population-level vulnerability to climate change. We found a rich assortment of responses to climate change across the range of balsam poplar, including high local, forward, and reverse offsets and migration distances in the eastern portion of the range and low offsets and migration distances in the center of the range. This suggests that eastern populations may face the greatest vulnerability to climate change compared to the rest of the range and greatest risk of future extirpation. More broadly, our work shows that, just as some climates and biological communities may disappear from the future landscape, and novel ones may emerge in their place^10^, the same concepts apply to the genetic composition of climatically adapted tree populations. The concepts of forward and reverse genetic offset provide a new way to consider population-level risk to future climate change that accounts for local adaptation and goes beyond the constraints of species-level predictions.

## MATERIALS AND METHODS

### Balsam poplar

Balsam poplar is a northern broad-leaved forest tree species that occurs over a large portion of the boreal region of Canada and the northern United States. The expansive range of *P*. *balsamifera* spans more than 30 degrees of latitude across multiple broad climatic gradients^22^, with the center of its range in the boreal region of Canada that is expected to see amongst the highest levels of future warming in North America^23^. Trees in the *Populus* genus have emerged as a model system for landscape genomic studies of local adaptation to climate^24^ and studies of balsam poplar, in particular, have shown it to be locally adapted to climate for numerous functional traits^20,25^ and its adaptive variation to be climatically structured^26^.

### Generalized dissimilarity models

We used generalized dissimilarity models to map predictions of local adaptation to climate (Fig. S1). Generalized dissimilarity models (GDM^27^) are a type of non-linear matrix regression that accounts for the curvilinear relationship between climatic (and optionally, geographic) distance and genetic differentiation among populations separated along environmental gradients. We fit GDM to genetic differentiation (*F*_*ST*_) of 33 single nucleotide polymorphisms (SNPs) in the *Populus* flowering time gene network genotyped in 995 individuals from 81 populations from across the range of balsam poplar^19^. Genes in the flowering time network are associated with both reproductive and vegetative plant phenology, by regulating the timing of seasonal growth, dormancy, and reproduction with the permissive growing season. We selected SNPs that had a relationship with environment, identified using Bayenv2^28^ and latent factor mixed models^29^. The environmental variables assessed for a relationship with SNPs included the first 3 components of a principal components analysis (PCA) of 19 bioclimatic variables and elevation, meant to capture the dominant climatic gradients across balsam poplar’s range. See Keller et al.^19^ for a full description of how candidate SNPs were identified. We used SNPs showing an association with any of the 3 PCA axes identified across a range-wide sample (see Table 2 in^19^). Based on these 33 climate-associated candidate SNPs, we calculated a multi-locus pairwise *F*_*ST*_ among the 81 populations using the ‘genet.dist’ function in the hierfstat package^30^ in R. Any pairwise *F*_*ST*_ values less than zero were assigned a value of zero.

GDM was parameterized with six bioclimatic variables that lacked strong correlation (|r| < 0.75). These included: summer and winter mean temperature (bio10, bio11) and precipitation (bio18, bio19), isothermality (bio3), and mean diurnal range (bio2) downloaded from the WorldClim dataset^31^, at a resolution of 10 arc-minutes. We tested for variable importance in the model by permuting each variable 100 times and determining the average decline in deviance explained after permutation. Models were parameterized in current climate (centered on ~1975) and predicted to future climate (centered on 2070) using a composite average of five global circulation models (GCMs; UCAR Community Climate System Model, NOAA Geophysical Fluid Dynamics Laboratory Coupled Physical Model, MET Office Hadley Center Earth System Model, NASA Goddard Institute for Space Studies-E2-R, and the Norwegian Earth System Model). We performed all analyses using two different emission scenarios (RCP 4.5 and 8.5) for 2070. Results and discussion refer to the composite mean of the five RCP 8.5 projections for 2070 unless specifically noted. GDMs were fit using the gdm package^32^ in R.

### Genetic exposure metrics

The GDM used to predict genetic exposure metrics explained 65.2% of the deviance in *F*_*ST*_. Variable permutation revealed that winter precipitation was the most important variable in the model. Isothermality and summer temperature were of secondary importance, while summer precipitation, mean diurnal range, and winter temperature were least important.

We used GDM to quantify the disruption of adaptive gene-climate associations expected under climate change using three different formulations of ‘genetic offset,’ which we term: (i) local, (ii) forward, and (iii) reverse genetic offset (Fig. 1). Following Fitzpatrick & Keller^8^, local offset (a metric of predicted maladaptation to future climate within a focal site) was calculated by predicting *F*_*ST*_ for locally adaptive SNPs between present and future climate at the same location, assuming no migration or gene flow. Forward offset is the minimum expected disruption in the gene-climate association assuming populations have unlimited dispersal capacity. Forward offset was quantified by first predicting *F*_*ST*_ between current climate at each focal grid cell within balsam poplar’s current range and all grid cells in North America (exclusive of Mexico) for future climate. From these predictions, we then identified the future climate grid cell with the minimum predicted *F*_*ST*_, which we term forward *F*_*STmin*_ or forward offset. The distance to the location that minimized forward offset represents the minimum migration distance required to minimize maladaptation (*D*_*min*_). High values of forward offset indicate maladaptation to all future North American climates, and indicate the possibility that contemporary gene-climate associations may disappear from the landscape. To assess the sensitivity of forward offset to dispersal constraints, we tested how *F*_*STmin*_ varied when migration was limited to five distance classes (50, 10, 250, 500, 1000 km). In addition to geographic distances, we also calculated the initial bearing populations would follow if they were to migrate to the location that minimized forward offset. Distance and bearing were calculated with the ‘distGeo’ and ‘bearing’ functions, respectively, in the geosphere package^33^ in R.

Reverse offset follows the same idea as forward offset, but is calculated from future climate to current climate. In this case, we first used the GDM (the same model discussed above) to predict *F*_*ST*_ between each future climate grid cell within balsam poplar’s current range and all current climate grid cells within balsam poplar’s current range. From these predictions, we then identified the current climate grid cell with the minimum predicted *F*_*ST*_, which we term reverse *F*_*STmin*_ or reverse offset. Reverse offset provides a metric of how novel the future gene-climate association is predicted to be at a given site, relative to existing gene-climate associations present at any location throughout balsam poplar’s range under current climate. As such, high values of reverse offset indicate “genetic novelty” as there is no analogous gene-climate association found anywhere in the current landscape. Note that for reverse offset we only considered pixels within balsam poplar’s current range at both times periods to ensure future novelty in gene-climate associations was quantified only with respect to locations where balsam poplar currently occurs (i.e., within the current range) as opposed to the entirety of North America as was done for forward offset.

To simultaneously visualize local, forward, and reverse offset, we mapped these three metrics as the red, green, and blue bands of an RGB image, respectively. Because values of local offset were systematically higher than forward or reverse offset, we rescaled values within each band to their quantiles before plotting. This ensured the full range of colors were possible in the RGB images, and is analogous to a histogram equalization performed on each band. We also tested for correlation between local, forward, reverse offsets and distances. We used a spatially-corrected Pearson’s correlation coefficient to quantify these relationships. The spatial correlations were implemented with the SpatialPack package^34^ in R, after projecting latitude/longitude coordinates to an equidistant projection (Azimuthal equidistant). Upon publication, R code to calculate local, forward, and reverse offsets will be available at github.com/agougher/poplarAdaptiveOffset.

---

**Fig. S1.**
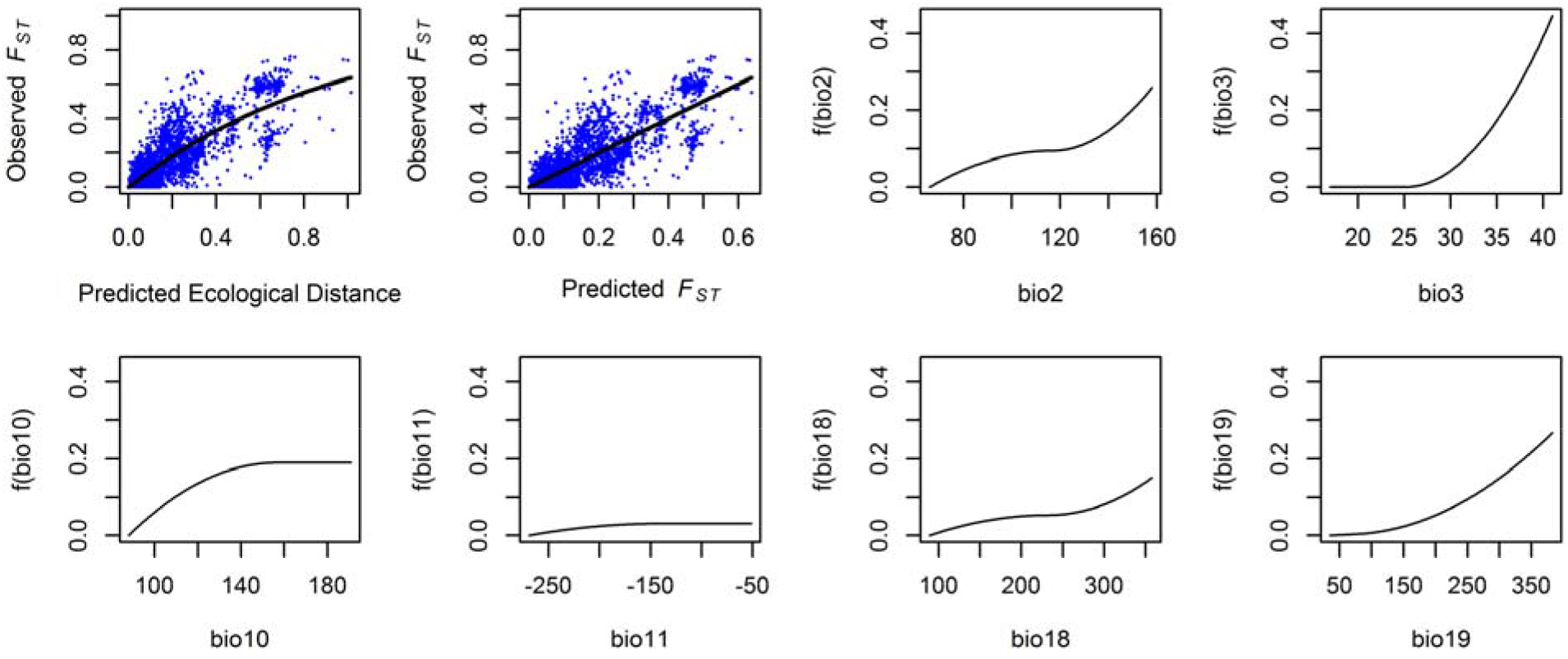
Generalized dissimilarity model (GDM) fit and climatic response plots. GDM was fit to *F*_*ST*_ of 33 SNPs in the *Populus* flowering time network across 81 range-wide balsam poplar populations (bio2: mean diurnal range; bio3: isothermality; bio10: mean summer temperature; bio11: mean winter temperature; bio18: summer precipitation; bio19: winter precipitation).

**Fig. S2.**
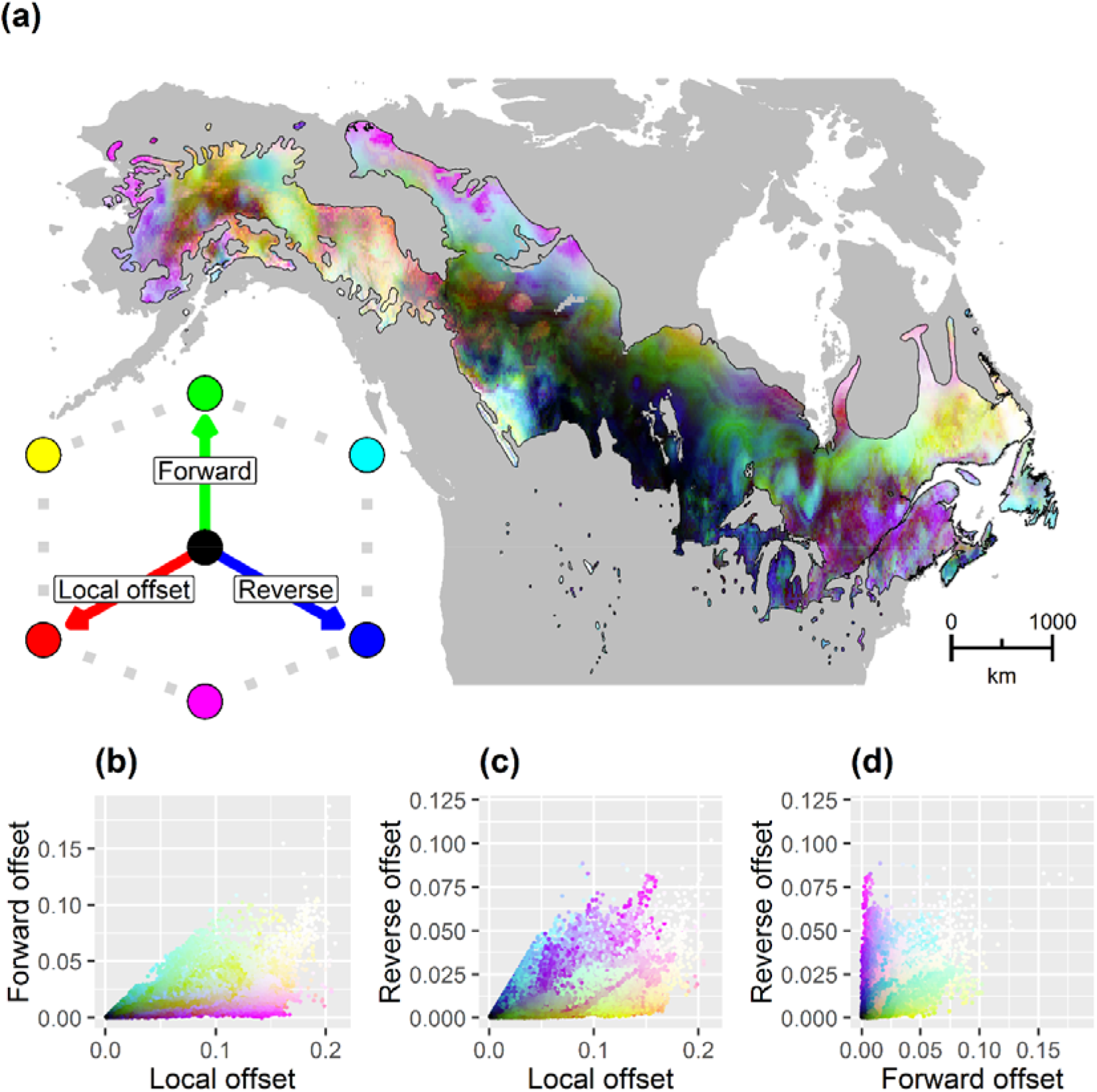
Red-green-blue map of local (red), forward (green), and reverse (blue) offset throughout the range of balsam poplar for 2070 and RCP 4.5. Brighter colors (closer to white) have relatively high values along each of the three axes indicating greater predicted exposure to climate change, while darker colors (closer to black) have relatively lower values, indicating lower exposure to climate change. (b-d) Bivariate scattergrams of (a). All units are *F*_*ST*_, but migration and novelty offsets are minimized *F*_*ST*_.

**Fig. S3.**
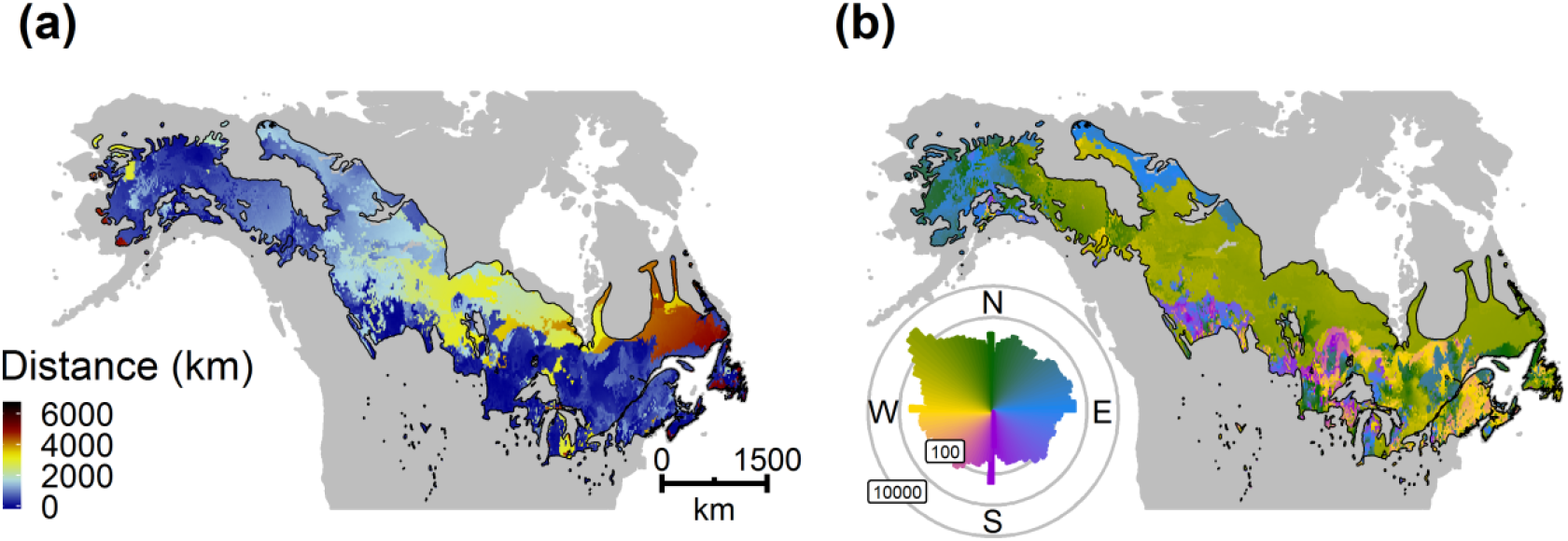
(a) Distance and (b) initial bearing to the location that minimizes future offset for balsam poplar in 2070 and RCP 4.5. Polar histogram in (b) shows the log10 number of cells in each bearing bin.

**Fig. S4.**
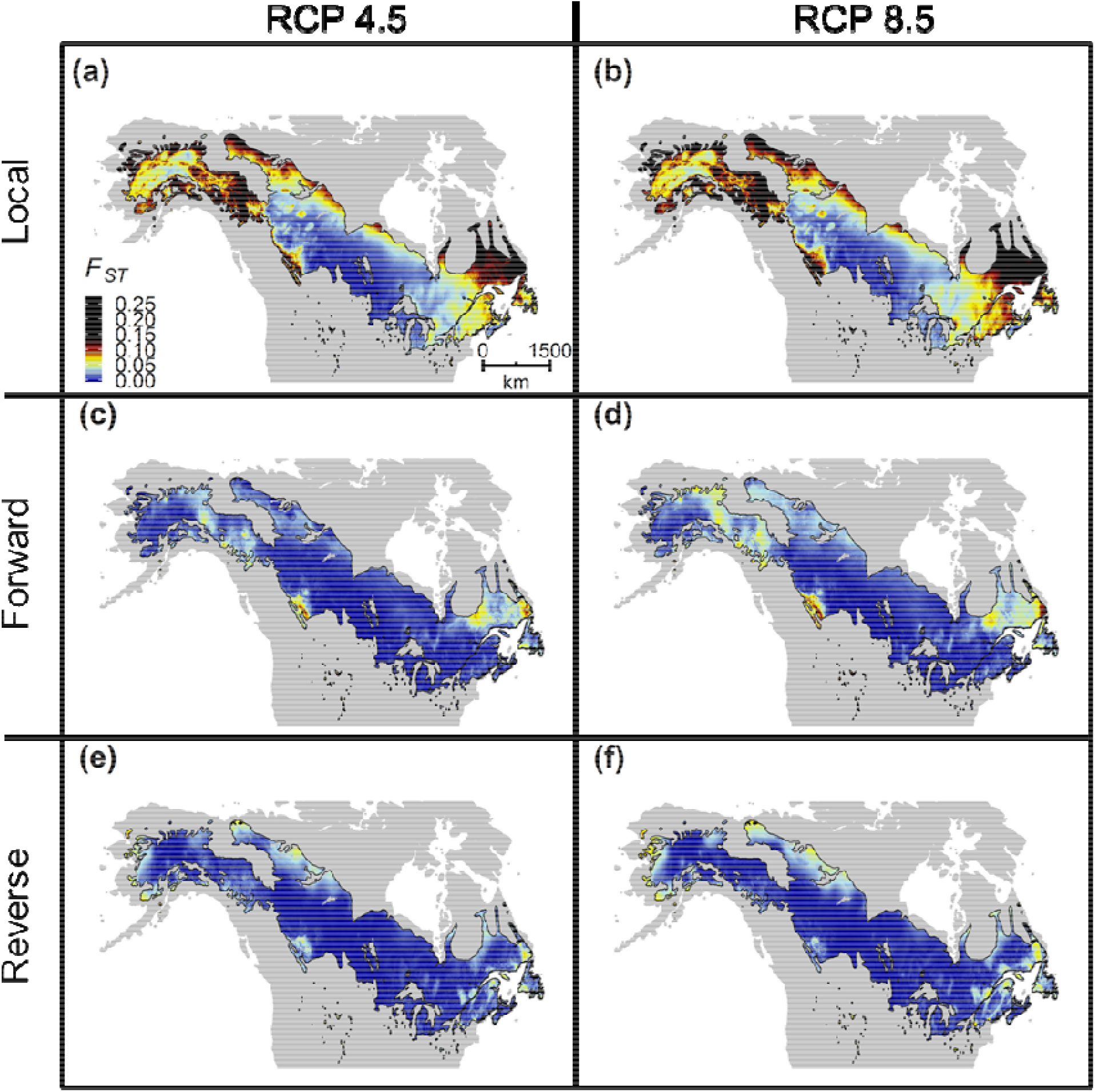
(a & b) Local genomic offset, (c & d) forward offset, and (e & f) reverse offset for RCP 4.5 (first column; a, c, e) and RCP 8.5 (second column; b, d, f) for 2070. Note the non-linear color scale. a, c, e are plotted as an RGB image in Fig. S2, and b, d, f are plotted in Fig. 2.

**Fig. S5.**
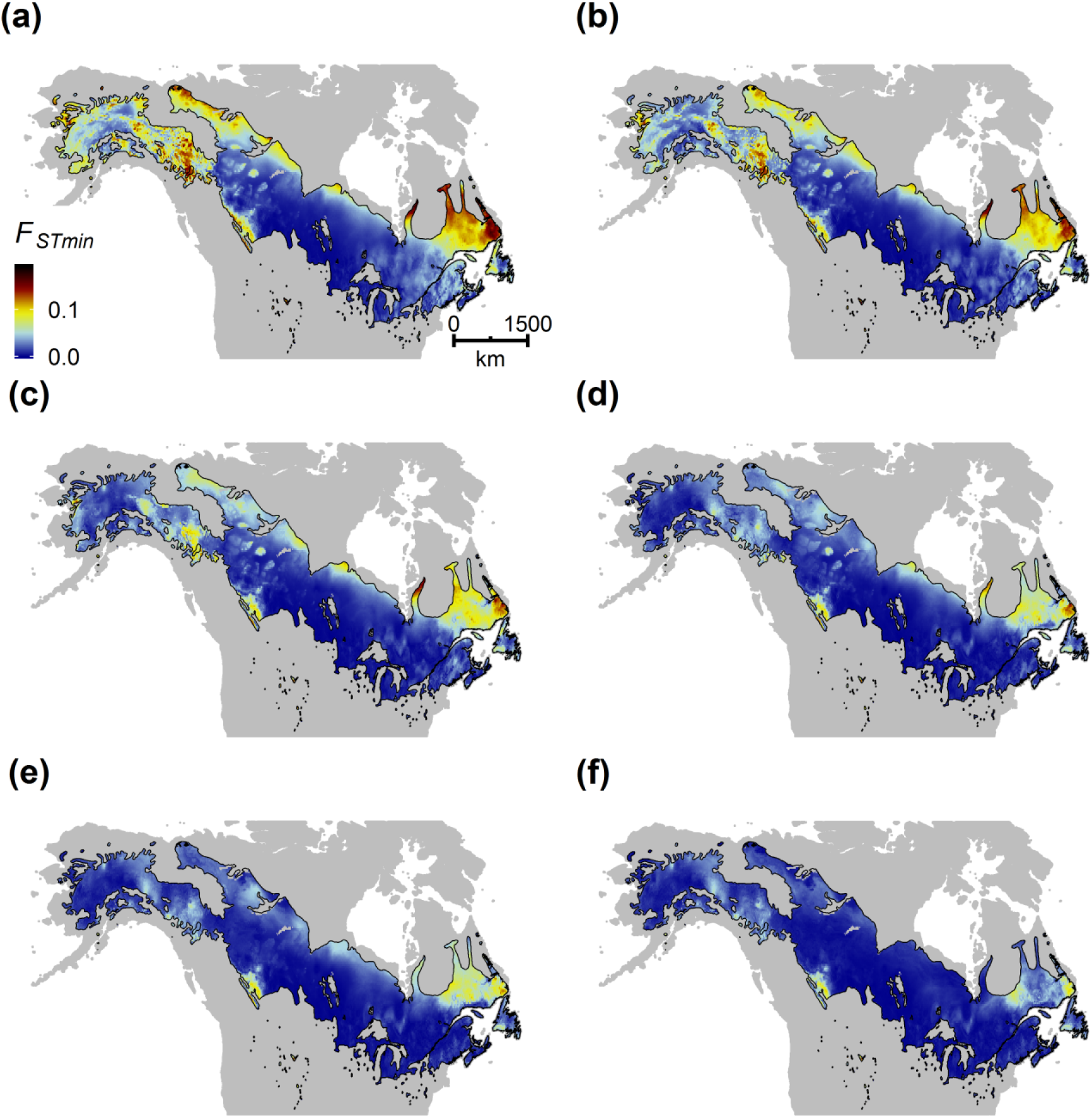
Effect of search distance on forward *F*_*STmin*_ for RCP 4.5 in 2070. Distance classes included (a) 50 km, (b) 100 km, (c) 250 km, (d) 500 km, (e) 1000 km, and (f) unlimited. See also Fig. 3.

**Fig. S6.**
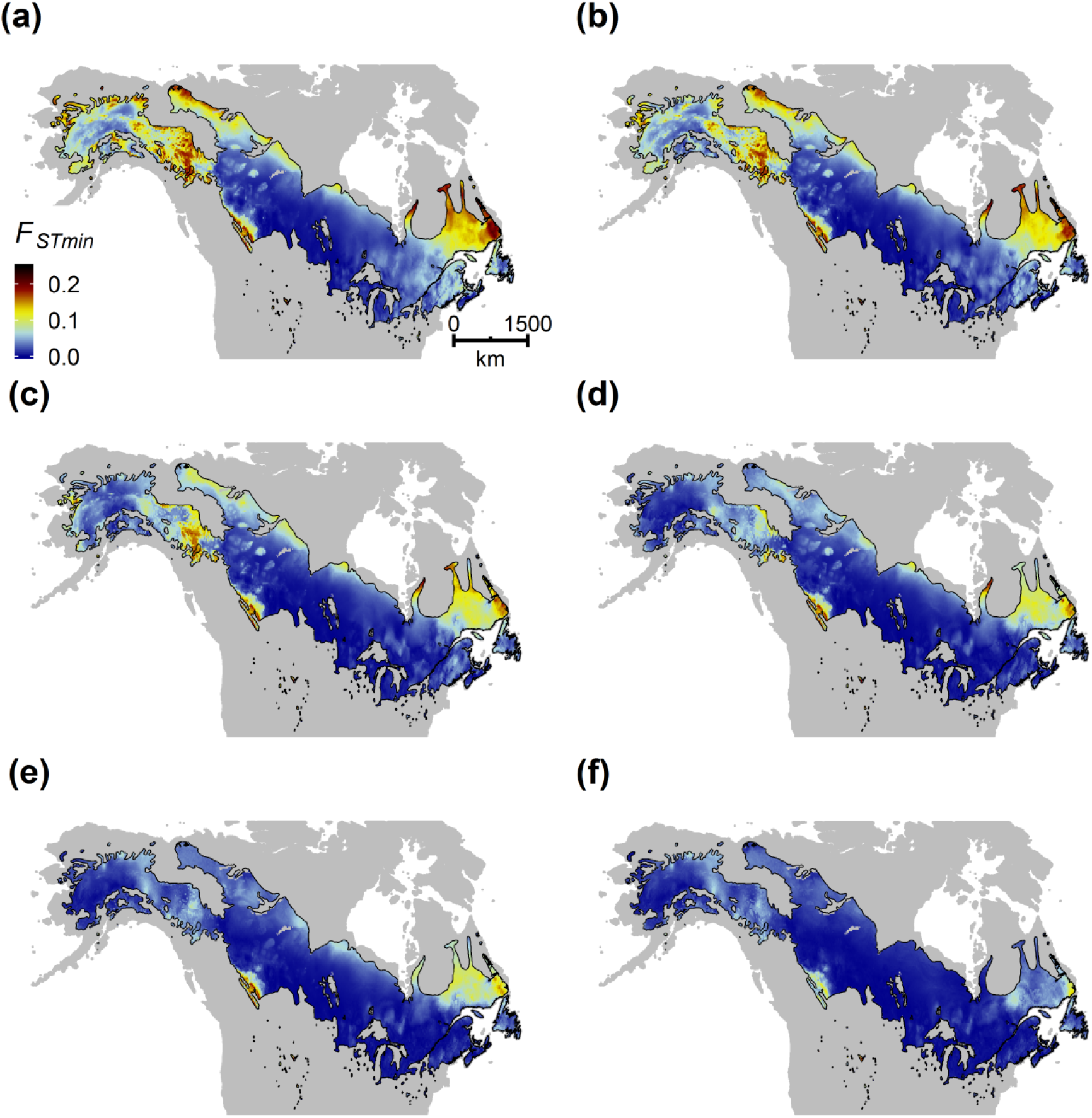
Effect of search distance on forward *F*_*STmin*_ for RCP 8.5 in 2070. Distance classes included (a) 50 km, (b) 100 km, (c) 250 km, (d) 500 km, (e) 1000 km, and (f) unlimited. See also Fig. 3.

